# Human-immune-system humanized-DRAGA mice are a valuable model to study novel immunotherapies for HIV-1

**DOI:** 10.1101/2024.12.02.626442

**Authors:** Pongthorn Pumtang-on, Negin Goodarzi, Brianna C Davey, Emily N Sevcik, Natalie Coleman-Fuller, Vaiva Vezys, Mangala Rao, Mary S Pampusch, Aaron K Rendahl, Sofia A Casares, Pamela J Skinner

## Abstract

Humanized (h)DRAGA mice are a promising *in vivo* model for investigating immunotherapies for treating HIV infections. These mice are not only susceptible to HIV infection, but they also develop functional human immune cells, including T cells and B cells as well as follicular-like structures that mimic lymphoid B cell follicles, where HIV-producing cells concentrate during infection in a manner similar to that found in humans. This study evaluated the safety, tissue targeting, and efficacy of follicular-targeting HIV-specific chimeric antigen receptor (CAR)-T cells (CAR/CXCR5-T cells) in HIV-infected hDRAGA mice. Intravenously-infused CAR/CXCR5-T cells persisted in hDRAGA mice for the duration of the study, peaking six days post-infusion. This study indicated that CAR/CXCR5-T cell treatment is safe, with 100% survival rate of treated mice and no noticeable changes in pathology. Six days after infusion, CAR/CXCR5-T cells had accumulated in the follicle-like structures, with many appearing in direct contact with HIV-producing cells. However, CAR/CXCR5-T cell treatment did not appear to reduce viral loads compared to controls, perhaps because many of the engineered CAR/CXCR5-T cells were themselves infected with HIV, with some CAR/CXCR5-T cells showing evidence of HIV virion release. Future studies will investigate whether CAR/CXCR5-T cells engineered for resistance against HIV infection are effective in reducing viral loads. This study supports the approach of using the HIV-infected hDRAGA mouse model to test cellular immunotherapies for HIV, as the model recapitulates many aspects of HIV infection in human lymphoid follicles.

## Introduction

In 2022, it was estimated that 39.0 million people were living with human immunodeficiency (HIV) worldwide, with 1.3 million people acquiring HIV infection during the year (1). Although antiretroviral therapy (ART) is effective in suppressing HIV, it is not a cure. Upon cessation of ART, viremia ensues. Thus, there is a need to create improved therapies for HIV infections that lead to sustained long-term virologic control.

Chimeric antigen receptor (CAR)-T cells are a promising therapy for targeting HIV, as they can be engineered to be directly cytotoxic to cells expressing specific viral targets on their surfaces (2). First introduced in the field of cancer research, CAR-T cells have been used to successfully treat B cell malignancies (3–6). We developed HIV- and SIV-specific CAR/CXCR5-T cells containing a bispecific CD4-mannose-binding lectin (MBL) CAR (7) and encoding a follicular homing receptor (CXCR5) to target B cell follicles (8, 9). HIV- and SIV-specific CAR/CXCR5-T cells showed specific cytotoxic functions and suppression *in vitro* (7–9). In an SIV-infected rhesus macaque (RM) model of HIV infection, CAR/CXCR5-T cells accumulated in lymphoid follicles, came in direct contact with SIV-producing cells, and in some instances, were associated with decreased SIV viral loads (10).

One major barrier for HIV therapeutic studies is the availability of animal models to study HIV pathogenesis. While non-human primate (NHP) models infected with SIV mimic many pathologies caused by HIV, the use of NHP models is complicated by their genetic diversity, particularly at immunogenetic loci, which can determine their susceptibility to and maintenance of sustained infection (11). Such inconsistencies in NHP models can impose limitations on developing HIV therapeutics, as different types of NHPs infected with various strains of SIV can show as much as 30% spontaneous control of infection to low or undetectable levels in plasma (12, 13), potentially complicating analyses of therapeutic efficacies. Additionally, most HIV immunotherapies cannot be used directly in SIV-infected NHP models of HIV because the HIV target molecules are not present in SIV. Given these limitations, various simian/HIV (SHIV) viruses have been investigated as models for HIV because they express HIV envelope (14–21). However, the inconsistency of viremia and the finding that many NHPs spontaneously control SHIV infections, make these models problematic for testing the efficacy of HIV immunotherapies (22–24). Consequently, findings from SIV/SHIV studies in NHP models may not directly translate to HIV therapeutic testing.

Humanized mice are valuable *in vivo* models for studying HIV pathogenesis (25, 26), HIV cellular immunotherapies (27–30), and HIV vaccines (31, 32), as they are susceptible to HIV infection, sustain high viral loads, show CD4^+^ T cell depletion, and maintain HIV reservoirs (33). Importantly, products developed for treating HIV infections can be tested directly in HIV-infected humanized mice, without modification. The development and use of HLA-transgenic humanized mice improve reconstitution of functional human T and B cells after infusing HLA-matched hu-HSC (34–36). Humanized (h) DRAGA mice have been engineered to express human HLA-A2.1 and HLA-DR0401 transgenes on a Rag1KO.IL2RγcKO.NOD (NRG) background (37). These mice robustly develop human B and T cells, B cell immunoglobulin class-switching, and elicit specific human cellular and antibody responses after vaccination (37–40). The hDRAGA mice are susceptible to infection with many human pathogens, including HIV (38, 39, 41–44). Unlike other humanized mice, hDRAGA mice develop follicle-like structures in lymph nodes and spleen (43), a critical feature of the model, as HIV replication is concentrated in lymphoid follicles (45–49). Importantly, HIV virions were shown to replicate in follicular-like CD20^high^ areas of secondary lymphoid tissues of HIV-infected hDRAGA mice (43), thus making this model particularly valuable to study HIV infections and pathogenesis.

Here, we evaluated the hDRAGA mouse model to test an HIV immunotherapy. We produced and infused human HIV-specific CAR/CXCR5-T cells to HIV-infected hDRAGA mice and evaluated the safety and efficacy of the immunotherapeutic cells to treat HIV. As predicted, the CAR/CXCR5-T cells accumulated in B cell-containing follicle-like structures (CD20^high^ areas) and were observed in direct contact with HIV RNA^+^ cells. The CAR/CXCR5-T cells administered intravenously were consistently detected in HIV-infected hDRAGA mice throughout the study. However, the CAR/CXCR5-T cells were insufficient to reduce HIV-1 viral loads in plasma and were unable to prevent CD4^+^ T cell loss post-infection, likely because the infused CD4+ CAR/CXCR5+ T cells and non-transduced cells in the product were susceptible to HIV infection. Our findings support the use of the hDRAGA mice as an appropriate model to test cellular immunotherapies aimed to provide long-term durable virologic control.

## Materials and methods

### hDRAGA mice and ethical considerations

All animal procedures reported herein were conducted under IACUC protocols approved by the Walter Reed Army Institute of Research/Naval Medical Research Command (WRAIR/NMRC) (22-28-AVAT) and the University of Minnesota (UMN) (2011-38633A) in compliance with the Animal Welfare Act and by the principles outlined in the “Guide for the Care and Use of Laboratory Animals,” Institute of Laboratory Animals Resources, National Research Council, National Academy Press, 2011.

DRAGA mice were bred and humanized at the Veterinary Service Program at WRAIR/NMRC. De-identified umbilical cord blood positive for HLA-A2.1 and HLA-DR0401 were commercially procured through the New York Blood Center (Long Island City, NY, USA (https://nybloodcenter.org/products-and-services/blood-products/research-products/). Briefly, the mice were irradiated (350 rads) and injected intravenously with CD3^+^ T cell-depleted cord blood (EasySep Human CD3 Positive Selection Kit, 18051, Stem Cell Technologies) containing approximately 10^5^ human CD34^+^ hematopoietic stem cells (HSC) as determined by FACS using a mouse anti-human CD34 antibody (BD Biosciences, 550761). CD3 depletion of human T cells was required to avoid lethal (acute) graft-versus-host-reaction. In this study, the mice were infused with hu-HSC from one human cord blood donor (HLA halotype A02:01A11:01B15:01B52:01DR04:01DR15:02), and the percentages of human T (CD3^+^, CD3^+^CD4^+^, and CD3^+^CD8^+^) and B (CD19^+^) cells were examined at WRAIR/NMRC at 18 weeks post-humanization. The procedures for assessing percentages of human T and B cells by FACS on the mononuclear FSC/SSC gate have been described previously (37). Following the infusion, the hDRAGA mice were transferred to UMN. Prior to HIV infection, the hDRAGA mice were reevaluated for human reconstitution status by assessing the percentages of human CD45^+^ and human T cells in peripheral blood by flow cytometry.

### Animal study, HIV-1 infection of hDRAGA mice, and experimental design

Fourteen female hDRAGA mice (designated as DRAGA no. 1-14) were housed in UMN animal facilities. Housing conditions were temperature-controlled (22°C ± 1°C) on a light/dark cycle of 14 h/10 h with *ad libitum* access to irradiated food and acidified water (pH 2.9 ± 0.2). In this study, the mice were allocated into two groups: 1) control animals (HIV-infected only; n=6) and 2) treated animals (HIV-infected, CAR/CXCR5-treated; n=6). The remaining two animals without HIV infection were used for CAR/CXCR5-T cell production. The hDRAGA mice were infected with HIV-1Ba-L as described previously (26, 50). Twelve hDRAGA mice were pre-treated subcutaneously with 2.5 mg per 50 μl of medroxyprogesterone acetate (DMPA) (Mylan Institutional LLC) 7 days prior to intravaginal infection with 10,000 tissue culture infectious dose 50 (TCID50) of HIV-1Ba-L in 20 μl PBS. HIV-1Ba-L viruses (ARP-510) were obtained from the NIH HIV reagent program (51, 52). The hDRAGA mice were checked biweekly for HIV-1 levels in plasma. The hDRAGA mice were checked biweekly for HIV-1 levels in plasma. The hDRAGA mice testing negative for HIV-1 at 2 weeks post-inoculation were reinfected with the same infectious dose of HIV and rechecked for HIV-1 levels in plasma. After the plasma HIV-1 viral loads had plateaued, the transduced CAR/CXCR5 T cell products were infused into the treated animals. The hDRAGA mice were sacrificed at 6, 14, and 28 days post-infusion (DPI).

### Production of CAR/CXCR5-T cells from the spleens of hDRAGA mice

Spleens from two non-HIV-infected hDRAGA mice were collected and disaggregated. Disaggregated cells from one of these mice were used to produce engineered CAR/CXCR5-T cells containing a bispecific CD4-MBL CAR and encoding CXCR5 by adapting our published methods (8–10, 53), as described previously (9, 53). The transduction efficacy of CAR/CXCR5-T cells was evaluated by assessing the co-expression of MBL and CXCL5 in live CD3^+^ cells using flow cytometry. The gating strategy included the following steps: identifying lymphocytes (SSC-A vs. FSC-A), singlets (FSC-H vs. FSC-Width), viable cells (SSC-A and Live/Dead NIR), mouse hematopoietic cells (SSC-A and mCD45^+^), human hematopoietic cells (SSC-A and hCD45^+^), human T cells (SSC-A and CD3^+^), and finally, CAR/CXCR5-T cells (MBL^+^ and CXCR5^+^).

### Infusion of transduced CAR/CXCR5 cell products

Prior to CAR/CXCR5-T cell infusion, the control and treated animals were allocated based on age, weight, peak of plasma HIV-1 RNA viral loads, levels of plasma HIV RNA viral loads before the infusion of transduced cell products, CD4^+^/CD8^+^ ratios, and CD4^+^ T cell count (**Supplemental Fig. 1**). The treated animals were infused intravenously with transduced cell products (2 × 10^5^ cells/g bodyweight). The animals were monitored twice daily for any signs of pain, illness, and stress by observing appetite, stool, behavior, and physical condition in response to the infused CAR/CXCR5-T cells.

### Blood and tissue collections

Peripheral blood samples were collected in blood collection tubes containing EDTA anticoagulant (Sarstedt AG & Co., 41.1395.105) pre-and-post infection, the day of CAR/CXCR5-T cell infusion, and at 6, 14, and 28 DPI. Peripheral blood (up to 100 μl) was centrifuged at 1,200 × *g* for 10 min, and plasma was collected as described previously (26, 43). The remaining peripheral blood (50 μl) was processed for cell phenotypes and CD4^+^ T cell count.

Spleens and lymph nodes were disaggregated using a 70 µm nylon cell strainer (Corning, 352350). For the spleens, red blood cells were lysed with ACK lysing buffer (Gibco, A1049201) according to the manufacturer’s instructions. Single-cell suspensions were stained for analysis using flow cytometry.

### Quantification of plasma HIV-1 viral loads

Cell-free HIV-1 RNA was isolated from 50 μl plasma using a QIAamp Viral RNA Mini Kit (QIAGEN, 52906) according to the manufacturer’s instructions. HIV-1 quantitation real-time PCR was performed using an HIV Quantitative TaqMan RT-PCR Detection Kit (Norgen Biotek Corp.) and a CFX96 Real-time PCR System (C100 Touch). Data were collected and analyzed with the CFX Manager 3.1 software (Bio-Rad, CA, USA).

### Quantitation of CD4^+^ T cell count in peripheral blood

Quantification of CD4^+^ cell numbers in peripheral blood was conducted using flow cytometric frequency gates, as described previously (54). Briefly, peripheral blood samples (50 μl) were treated with ACK lysing buffer (Gibco, A1049201) to lyse red blood cells. After centrifugation, cells were resuspended in 200 μl 1× PBS; 50 μl of the cell suspension was mixed with the AccuCheck counting beads (Thermo Fisher Scientific, PCB100) and counted on a CytoFLEX flow cytometer (Beckman Coulter), while the remaining suspended cells were stained for cell phenotyping. The data were analyzed with FlowJo software version 10 (BD Life Sciences). The CD4^+^ T cell count was analyzed based on the percentage of hCD3^+^MBL^-^CD4^+^ in lymphocytes compared to the number of beads and lymphocytes.

### Flow cytometry and antibodies

The resuspended cells were stained with anti-mouse antibodies mCD45 (BioLegend, 30-F11, 103149) and the following anti-human antibodies: hCD45 (BD Pharmingen, HI30, 557748), CD4 (BD Pharmingen, M-T477, 556615), CD3 (BD Pharmingen, P34.2, 557917), Live/Dead NIR (Invitrogen, L34976A), CD185 (CXCR5) (MU5UBEE, 12-9185-42), MBL (Invitrogen, 3E7, custom-made and labeled with Alexa Fluor 647, Invitrogen, A20186); and CD8 (BioLegend: SK1, 344731). Flow cytometric data were acquired on a Beckman Coulter CytoFLEX and analyzed with FlowJo software version 10 (BD Life Sciences).

### RNAscope in situ hybridization combined with immunofluorescence

In this study, we selected three slides (10 slices apart) from one treated hDRAGA at 6 DPI and one slide from one control hDRAGA at 6 DPI. RNAscope multiplex fluorescent kit V2 (Advanced Cell Diagnostics, UM 323100) with the opal fluorophores system (Akoya Bioscience) were used to simultaneously detect HIV vRNA^+^ and transduced CAR/CXCR5 cells according to the manufacturer’s instructions and as described previously (10, 55). For duplex detection of HIV vRNA+ and transduced CAR/CXCR5 cells, the sectioned slides were incubated overnight at 40°C with the C2 probe (Advanced Cell Diagnostics, V-HIV-1-Clade B 416111-C2) to detect HIV vRNA^+^ cells and a customized C1 probe to detect the CAR/CXCR5 cells. The sections were then washed with 0.5× RNAscope wash buffer and incubated with amplification reagents (1–3) according to the manufacturer’s instructions. Opal 570 and Opal 650 were used for the C1 and C2 probes, respectively.

For immunofluorescence staining, the sections were conducted as described previously (10) with some modifications. Briefly, the sections were stripped, washed, blocked, and incubated overnight at 4°C with 0.51 μg/ml rabbit anti-human CD20 antibodies (Abcam, EP459Y, ab78237) in 10% NGS-TBS-1% BSA. Then sections were washed and incubated with 7.5 μg/ml secondary AlexaFluor 488 conjugated polyclonal goat anti-rabbit IgG antibodies (Jackson ImmunoResearch, 111-545-003) in 10% NGS-TBS-1% BSA, counterstained with 1 μg/ml DAPI solution (Thermo Scientific, 2247) and mounted in Prolong Gold antifade reagent (Invitrogen, P36934). The stained tissue sections were imaged with a Leica TCS SPE DM6000 confocal microscope.

### Statistical analysis

All statistical analyses were performed using GraphPad Prism 10. Data are represented as mean ± SD, median, or geometric mean with 95% CI. Groups were compared using the Mann-Whitney test. A p-value of p < 0.05 was considered significant.

## Results

### hDRAGA mice reconstituted with human immune cells and infected with HIV showed sustained high HIV plasma viral loads

All 14 hDRAGA mice were assessed for human reconstitution at 18 weeks post-hu-HSC infusion at WRAIR/NMRC before the animals were transferred to UMN. The human immune cell reconstitution status in peripheral blood is shown in **Table I**.

**Table I.** Reconstitution of human immune cells in hDRAGA mice from WRAIR/NMRC.

| Mouse # | Frequency (%) of lymphocytes in peripheral blood |  |  |  |
| --- | --- | --- | --- | --- |
|  | CD19<sup>+</sup> | CD3<sup>+</sup> | CD3<sup>+</sup> CD4<sup>+</sup> | CD3<sup>+</sup> CD8<sup>+</sup> |
| DRAGA-1 | 57.7 | 8.5 | 7.9 | 0.6 |
| DRAGA-2 | 52.6 | 18 | 15.3 | 2.7 |
| DRAGA-3 | 75.8 | 10.8 | 8.7 | 2.1 |
| DRAGA-4 | 46.3 | 20.6 | 18.9 | 1.7 |
| DRAGA-5 | 64.2 | 13.3 | 11.8 | 1.5 |
| DRAGA-6 | 66.1 | 12.1 | 11.3 | 0.8 |
| DRAGA-7 | 61.7 | 12.2 | 10.7 | 1.5 |
| DRAGA-8 | 67.8 | 9.8 | 7.7 | 2.1 |
| DRAGA-9 | 69.0 | 9.4 | 7.4 | 2.0 |
| DRAGA-10 | 66.7 | 11.5 | 10.4 | 1.1 |
| DRAGA-11 | 61.0 | 9.7 | 9.1 | 0.6 |
| DRAGA-12 | 50.0 | 13.0 | 12.1 | 0.9 |
| DRAGA-13 | 54.6 | 10.6 | 9.7 | 0.9 |
| DRAGA-14 | 52.9 | 16.0 | 14.9 | 1.1 |

We re-examined the reconstitution of human immune cells in all hDRAGA mice (n=14) at 25 weeks after the hu-HSC infusion (**Fig. 1).** The 14 hDRAGA mice showed high frequencies of human hematopoietic-derived cells (**Fig. 1A**), identified by human CD45^+^ cells (63.51 ± 13.49%) in their peripheral blood samples (**Fig. 1B**). The hDRAGA mice showed a reconstituted profile of human CD3^+^ T cells (29.94 ± 8.51%), human CD4^+^ T cells (24.86 ± 6.7%), and human CD8^+^ T cells (2.69 ± 1.33%) (**Fig. 1C**).

**Figure 1.**
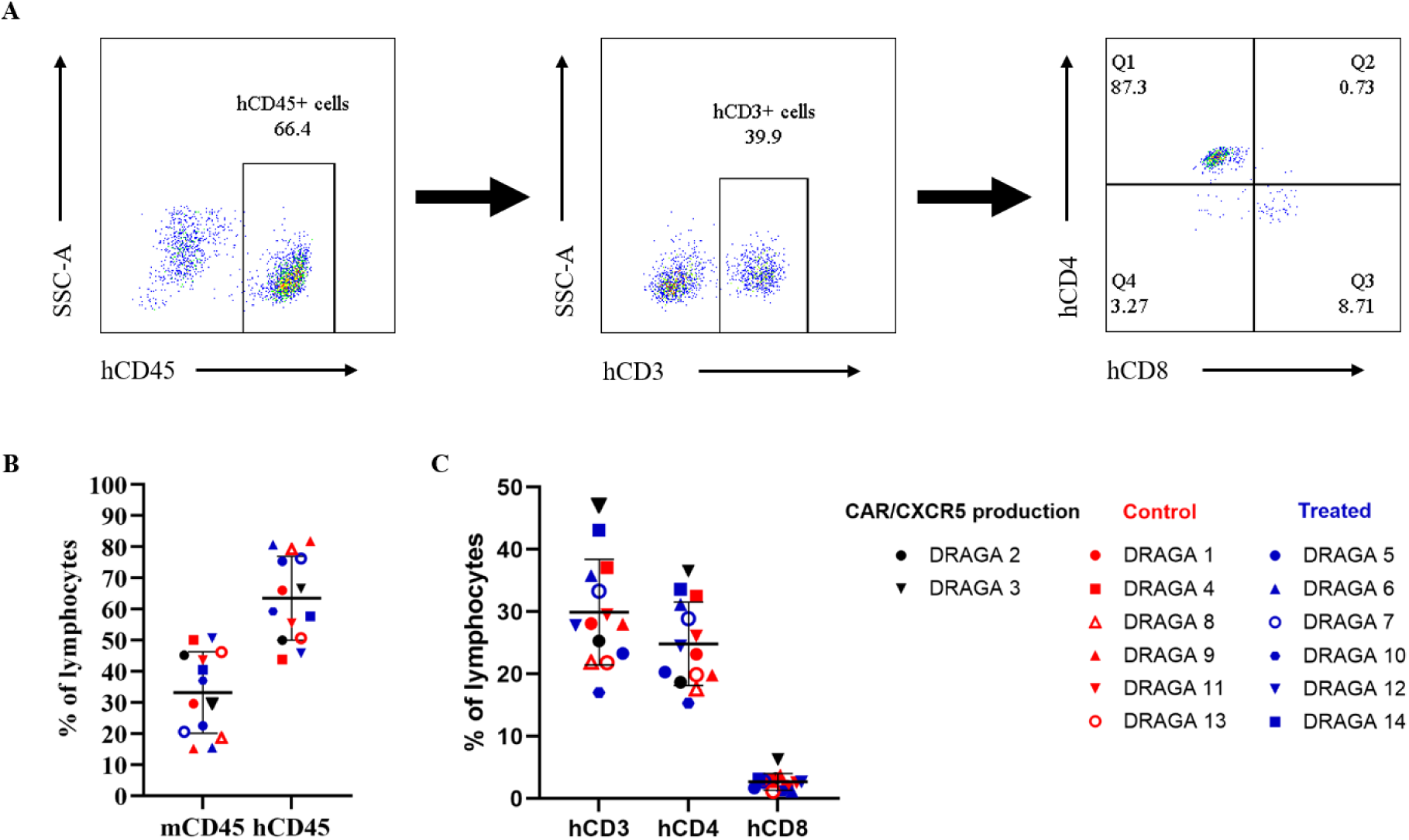
Human T cell reconstitution in peripheral blood from hDRAGA mice. Peripheral blood samples from all hDRAGA mice (n=14) were examined by flow cytometry for the level of human immune cell reconstitution prior to infection with HIV-1. (**A**) An example of the gating strategy for quantifying human immune cell reconstitution in a hDRAGA mouse. (**B**) Percentages of mouse CD45^+^ and human CD45^+^ cells in lymphocytes. (**C**) Percentages of human CD3^+^, CD4^+^, and CD8^+^ T cells. Data are mean ± SD.

Twelve hDRAGA mice were pretreated with DMPA and then infected with HIV-1. Six HIV-infected hDRAGA mice were treated with CAR/CXCR5-T cells and subsequently sacrificed to evaluate the efficacy and persistence of the CAR/CXCR5-T cells, as outlined in the experimental timeline for CAR/CXCR5 immunotherapy shown in **Figure 2**.

**Figure 2.**
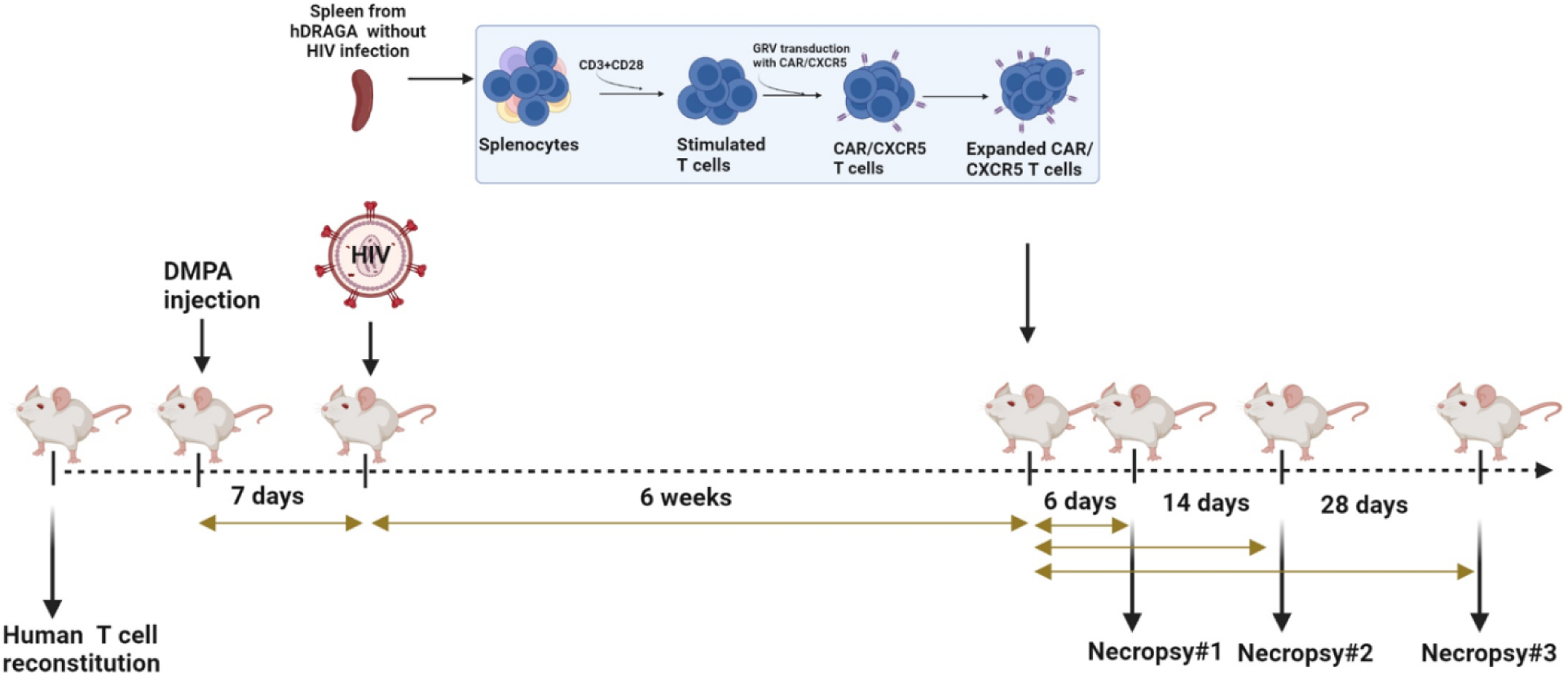
Timeline of CAR/CXCR5-T cell immunotherapy in hDRAGA mice. Animals (n=14) were re-evaluated for human cell levels, treated with DMPA (n=12), and 7 days later infected with HIV. Splenocytes from one of two uninfected hDRAGA mice were used to engineer CAR/CXCR5-T cells. Six weeks after HIV infection, animals were infused with CAR/CXCR5-T cell products or were untreated, with subsets of animals sacrificed at 6, 14, and 28 days post-treatment (DPT).

The hDRAGA mice were monitored biweekly for plasma HIV-1 viral loads after day 1 of HIV-1 inoculation, as assessed by qRT-PCR assays. Overall, 11 of the 12 (91.67%) hDRAGA mice were HIV-infected. Ten hDRAGA mice were positive for HIV-1 in their plasma by 4 weeks post-infection (wpi) with a single HIV-1 infection (**Fig. 3**). Two hDRAGA mice (DRAGA 8 and 11) were negative for HIV-1 infection at 4 weeks and were reinfected with HIV-1 (**Fig. 3**). Two weeks later, DRAGA 11 showed HIV infection, while DRAGA 8 remained below the detection limit, resulting in the exclusion of this animal in this study. The time of peak plasma viral load varied among the HIV-infected mice (**Fig. 3**).

**Figure 3.**
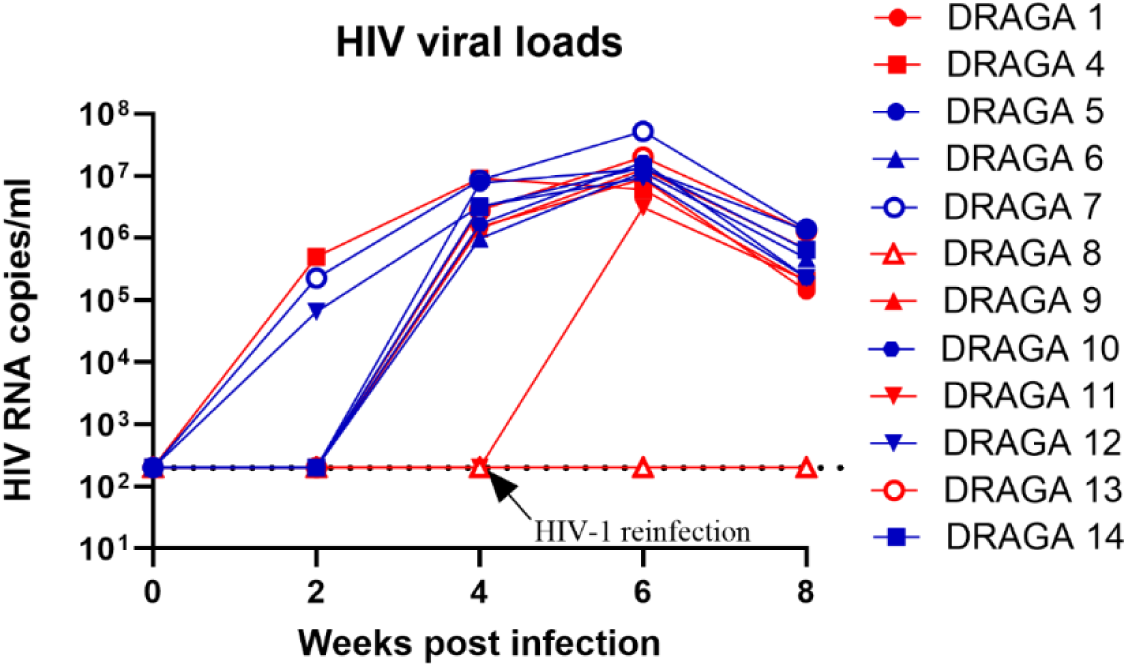
HIV-1 RNA plasma viral loads prior to CAR/CXCR5-T cell infusion. Peripheral blood samples were collected biweekly from hDRAGA via facial vein. DRAGA 8 and 11 were reinfected with HIV-1 at 4 weeks post-infection (wpi). The level of HIV-1 RNA in plasma was quantified using qRT-PCR. The threshold of HIV RNA detection was 200 copies/ml (dashed line).

Prior to infusion with CAR/CXCR5-T cells, 11 HIV-infected hDRAGA mice were allocated based on the criteria described in **Supplemental Fig. 1** which included age, the peak of plasma HIV RNA viral loads, levels of plasma HIV RNA viral loads prior to infusion of transduced T cell products, weight, human CD4+/CD8+ T cell ratios, and human CD4+ T cell counts. The average CD4+ T cell counts in the control animals (geometric mean 666, 95% CI: 259-1712 cells/μl) were lower than those in the treated animals (geometric mean 1590, 95% CI: 630-4009 cells/μl), but this difference was not significant (p = 0.093).

### Successful production of CAR/CXCR5-T cells from hDRAGA splenocytes

Disaggregated splenocytes from a non-HIV-infected hDRAGA mouse were used to produce CAR/CXCR5-T cells. Flow cytometric analysis showed that approximately 72% of CD3^+^ T cells co-expressed MBL (a portion of the extracellular domain of this CAR molecule) and CXCR5 molecules (**Fig. 4A**). Of the transduced cells expressing CAR/CXCR5, 72.5% were CD4^+^CD8^-^ and 27.5% were CD4^+^CD8^+^ T cells (**Fig. 4B**). Note that the CD4 antibodies also detect the CD4-MBL-CAR molecule and, in this CD4+CD8+ population, we are likely detecting CD8 T cells expressing the CAR molecule.

**Figure 4.**
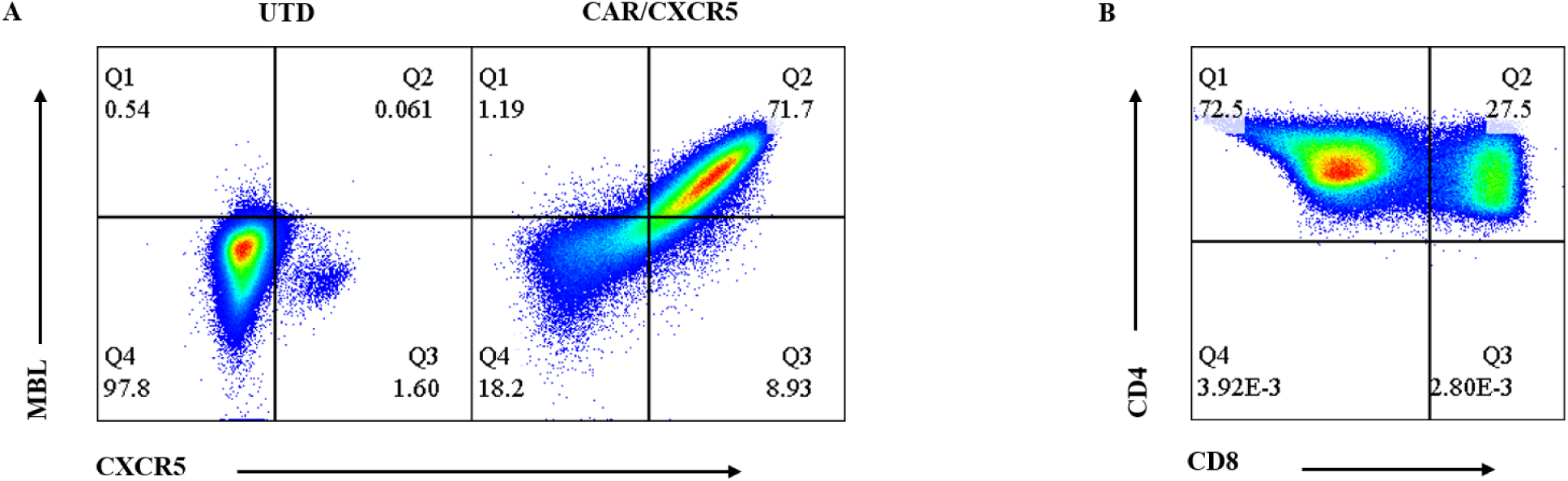
Characterization of transduced CAR/CXCR5-T cells produced from splenocytes of a hDRAGA mouse. (**A**) Anti-MBL was used to detect the CAR plots in untransduced (UTD) T cells (left) and transduced CAR/CXCR5-T cells (right). (**B**) CD4^+^ and CD8^+^ expression within CD4^+^MBL^+^CXCR5^+^ transduced T cells. Gates were set on lymphocytes, singlets, live human CD3^+^ T cells for A, and additionally on MBL^+^CXCR5^+^ cells for panel B. Cells expressing MBL and CXCR5 were considered to be CAR^+^.

We evaluated the safety of the CAR/CXCR5-T cell immunotherapy treatment in HIV-infected hDRAGA mice, compared to control mice after the infusion of the transduced CAR/CXCR5-T cell product. The animals exhibited no noticeable adverse effects after receiving the immunotherapeutic cells, and their weights were unaffected by the immunotherapeutic infusion. Necropsy showed no abnormalities of internal organs in treated animals beyond those typical in HIV-infected hDRAGA animals.

### CAR/CXCR5-T cells persisted in peripheral blood and secondary lymphoid tissues

After infusion, CAR/CXCR5-T cells were detected in the peripheral blood (**Fig. 5A**), lymph nodes (**Fig. 5B**), and spleens (**Fig. 5C**) of the treated mice through the end of the study at 28 DPI. In all the disaggregated cell samples, the number of CAR/CXCR5-T cells was highest at 6 DPI and gradually decreased over time.

**Figure 5.**
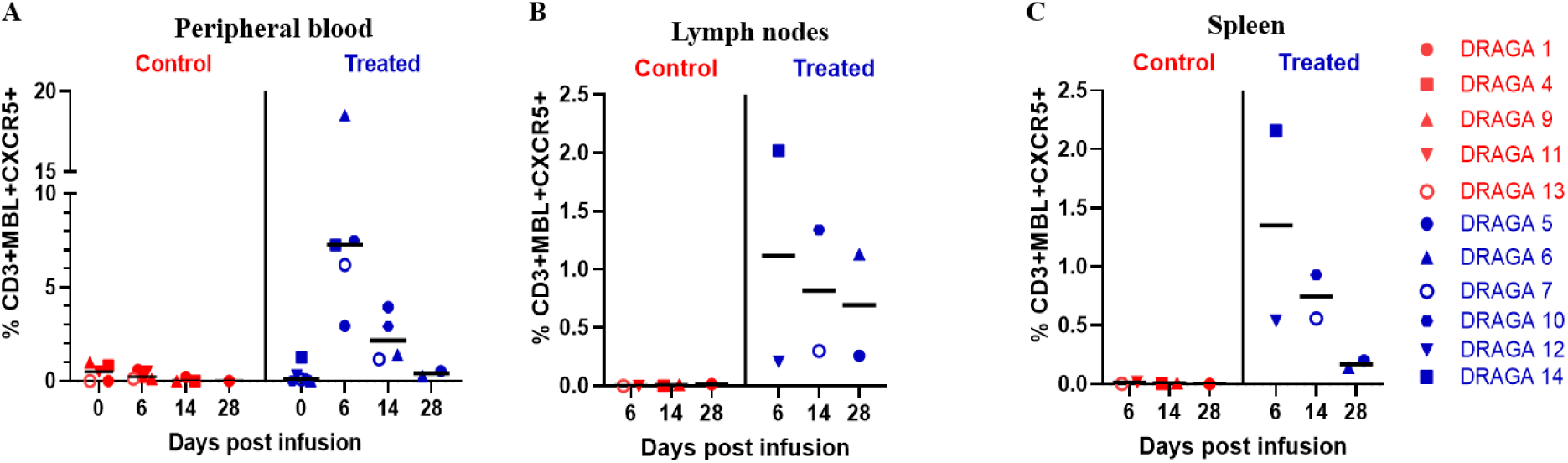
Persistence of CAR/CXCR5-T cells in HIV-infected hDRAGA mice post-infusion. CAR/CXCR5-T cells were enumerated in (**A**) peripheral blood, (**B**) lymph nodes, and (**C**) spleen of hDRAGA mice at 6, 14, and 28 DPI. Each graph shows the percentages of MBL^+^ and CXCR5^+^ (CAR^+^ T cells) cells within the live CD3^+^ T cell population. The bar represents median percentages.

### Transduced CAR/CXCR5-T cells were insufficient to reduce plasma HIV-1 viral loads

Our previous study in NHPs showed that infusion of CAR/CXCR5-T cells at a dosage of 2 × 10^8^ cells/kg body weight (equivalent to 2 × 10^5^ cells/g in mice) was safe and effective in reducing SIV viral loads in SIV-infected ART-treated and -released RMs (10). Thus, the same dosage of CAR/CXCR5-T cells was applied, although the hDRAGA mice were not ART-treated.

The difference in plasma HIV viral loads between treated and control animals was not statistically significant (p > 0.05 at all time points) (**Fig. 6A** and **Supplementary Table I**). The mean of CD4^+^ T cell counts slightly declined in both control and treated animals, but there was no statistically significant difference between the two groups at each time point (p > 0.05 at all time points) (**Fig. 6B**). Likewise, infusion of the CAR/CXCR5-T cell product did not impact the CD4+/CD8+ ratios (**Fig. 6C**).

**Figure 6.**
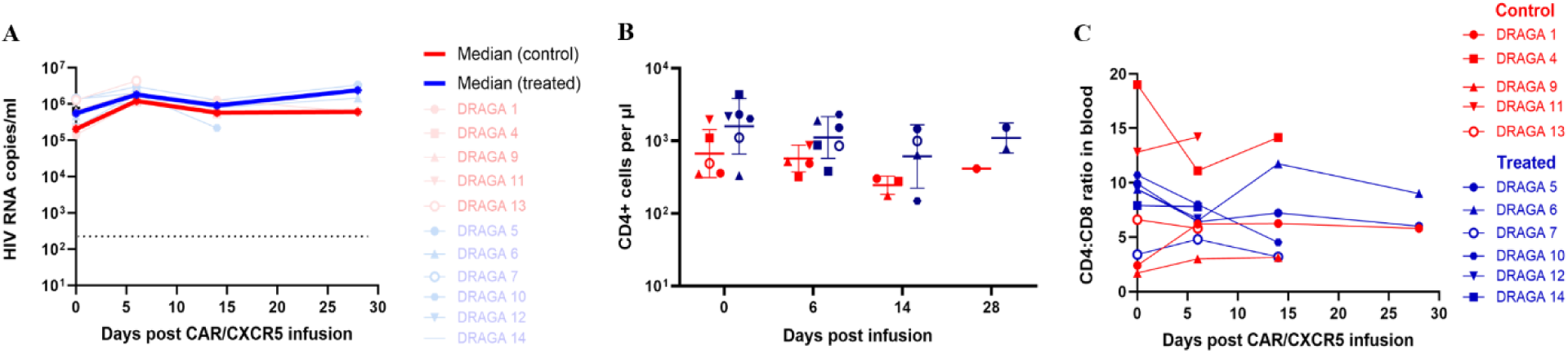
Effect of CAR/CXCR5-T cell infusion on plasma HIV viral loads and CD4^+^ T cell count. (**A**) The HIV viral loads were measured in the CAR/CXCR5-T cell-treated and -untreated control mice at 0, 6, 14, and 28 DPI using qRT-PCR. Differences were evaluated between control and treated animals using Mann-Whitney test (0 DPI, p = 0.833; 6 DPI, p = 0.833; 14 DPI, p > 0.99). The threshold of HIV RNA detection was 200 copies/ml (dashed line). (**B**) CD4^+^ T cell counts were conducted for the CAR/CXCR5-T cell-treated and -untreated control mice at 0, 6, 14, and 28 DPI. Differences were evaluated between control and treated animals using Mann-Whitney test (0 DPI, p = 0.126; 6 DPI, p = 0.310; 14 DPI, p =0.400). (**C**) CD4/CD8 ratios post CAR/CXCR5 cell infusion in the PBMCs. Red dots and blue dots represent the control and treated animals, respectively. The red and blue lines (A) represent the median viral loads in the control and treated animals, respectively. Bar and error (B) indicate mean ± SD. The red and blue lines with symbols (C) represent individual animals from each group.

### CAR/CXCR5-T cells were HIV vRNA^+^ within CD20^high^ areas

We investigated the localization of the CAR/CXCR5-T cells and HIV vRNA^+^ cells in a spleen of an infused hDRAGA mouse using RNAscope combined with immunohistochemistry (**Fig. 7A**). At 6 DPI, the spleen from a CAR/CXCR5-T cell-treated hDRAGA mouse (DRAGA no. 14) showed abundant accumulation of both HIV vRNA^+^ cells and CAR/CXCR5-T cells within the CD20^high^ B cell follicle-like structures (**Fig. 7A**). Importantly, many CAR/CXCR5-T cells were found in direct contact with HIV vRNA^+^ cells (**Fig. 7A**). However, as seen in Fig. 7A, many CAR/CXCR5-T cells were also HIV vRNA^+^. In a control HIV-infected hDRAGA mouse, HIV vRNA^+^ cells were detected and accumulated within the CD20^high^ B cell-containing follicle-like structures, and no CAR probe signal was detected (**Fig. 7B**).

**Figure 7.**
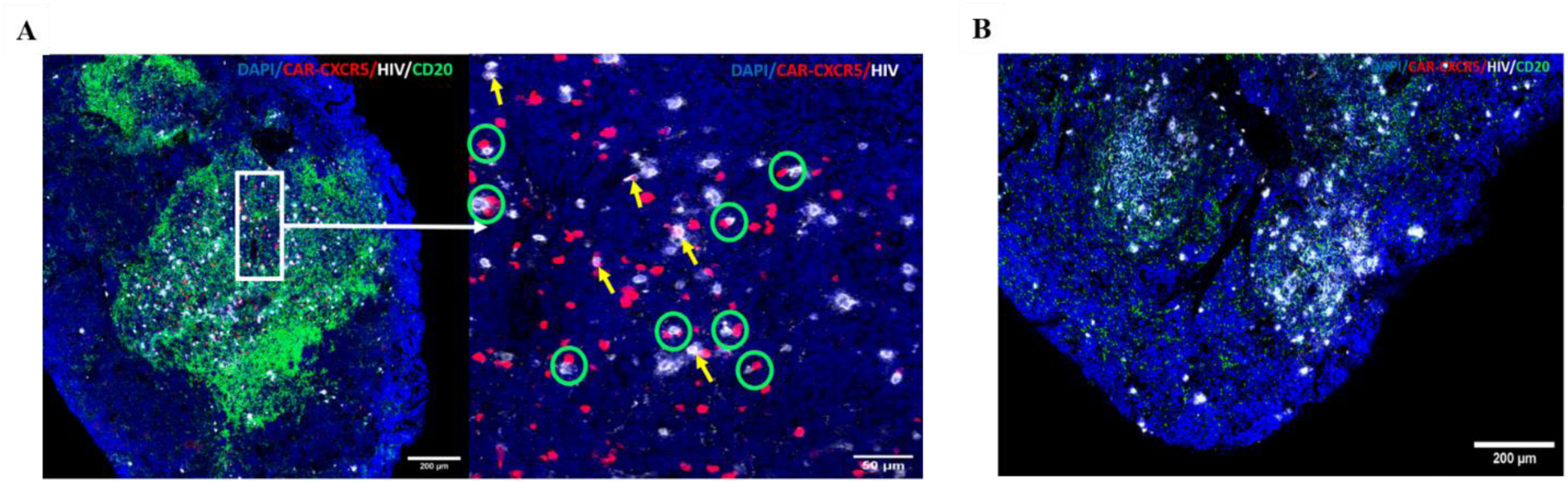
Localization of CAR/CXCR5-T cells and HIV vRNA^+^ cells. RNAscope in situ hybridization combined with immunohistochemistry showed that CAR/CXCR5-T cells (red) and HIV vRNA^+^ cells (white) localize in follicle-like structures of the spleen of a representative CAR/CXCR5-T cell-treated hDRAGA mouse and -untreated control HIV-infected hDRAGA mouse at 6 DPI. (**A**) In a spleen from a representative treated hDRAGA mouse (left), the enlarged image (right) shows CAR/CXCR5-T cells in direct contact with HIV vRNA^+^ cells, indicated in the green circles. Several CAR/CXCR5-T cells were HIV vRNA^+^ (yellow arrow). (**B**) RNAscope analysis of the spleen from a representative control HIV-infected hDRAGA mouse detected HIV vRNA^+^ cells (white) but not CAR/CXCR5-T cells (red). B cell follicle-like structures were identified using anti-human CD20 antibodies (green) and nuclei were stained with DAPI (blue). The confocal image was collected with a 20× magnification (NA = 0.4).

## Discussion

The current study demonstrated that hDRAGA mice had high levels of reconstituted human immune cells, were susceptible to HIV infection, and sustained high HIV-1 viral loads, supporting the findings of a recent study (43). We found that the HIV infection rate via intravaginal inoculation was more than 91%, which is similar to other studies that reported 100% HIV infectivity (26, 56–58). Our previous study showed that human HIV-specific (CD4-MBL) CAR/CXCR5-T cells were functional and suppressed HIV-infected cells *in vitro* (9). In this study, we investigated the safety and efficacy of human HIV-specific, follicle-targeting CAR/CXCR5-T cells in the hDRAGA mouse model.

Consistent with a previous study (43), we confirmed that the hDRAGA mice developed B cell-containing follicle-like structures in their secondary lymphoid tissues and HIV vRNA+ cells accumulated in these structures. In this study, we used the terminology of B cell-containing follicle-like structures instead of B cell follicles because the hDRAGA mice exhibit an atypical morphology of B cell follicle structures, such as a lack of canonical germinal centers and human follicular dendritic cells (43). We successfully produced CAR/CXCR5-T cells from the spleen of a hDRAGA mouse. The infused CAR/CXCR5-T cells persisted in follicular-like structures of the spleen through 28 DPI. These findings are consistent with our previous study investigating SIV-specific CAR/CXCR5-T cells infused in RMs (10). Furthermore, we demonstrated that the transduced CAR/CXCR5-T and HIV vRNA^+^ cells accumulated within CD20^high^ areas of the spleen, with many CAR/CXCR5-T cells in direct contact with HIV vRNA^+^ cells. These results showed that the transduced CAR/CXCR5-T cells effectively migrate to the B cell-containing follicle-like structures in secondary lymphoid tissues and come into direct contact with HIV-producing cells, indicating that they recognize HIV-producing cells.

However, many CAR/CXCR5-T cells became HIV vRNA^+^, indicating that the CAR/CXCR5-T cells were susceptible to HIV infection and likely contributed to the spread of infection. It is perhaps unsurprising that CD4^+^ CAR/CXCR5-T cells are susceptible to HIV infection due to the expression of HIV co-receptors CD4, CCR5, and CXCR4, given that our CAR/CXCR5-T cell product is composed of both CD4^+^ and CD8^+^ T cells. Our findings in the current study exemplify the need to engineer HIV-targeting CAR-T cells that are resistant to infection. Indeed, in studies by Maldini et al (29, 59), it was only after HIV-targeting CAR-T cells were made resistant to infection that differences in HIV viral loads were detected in treated versus control animals.

CAR/CXCR5-T cell treatment in HIV-infected hDRAGA mice failed to decrease the viral load, compared to untreated HIV-infected control mice, likely due to the fact that many CAR/CXCR5-T cells became infected and contributed to the spread of HIV infection, as described above. Of the CAR/CXCR5 T cell product, 72% were CD4^+^ T cells. In addition, approximately 30% of the infused cells in the product were non-transduced. Given that these cells are also targets of HIV infection, they likely also contributed to increased infection in the treated mice. Furthermore, this study was performed in chronically HIV-infected mice that were not ART-suppressed. Our previous study in SIV-infected ART-suppressed RMs detected very few CAR/CXCR5-T cells that were SIV vRNA^+^ (10). This study, which showed decreased viral loads in CAR/CXCR5-T cell-treated RMs compared to controls, was performed using ART-interrupted animals (10) where there were fewer virus-producing cells to clear relative to this study. Another possible reason the CAR/CXCR5-T cells were ineffective in this study at controlling HIV viremia could be that they were exhausted from the proliferation during uncontrolled HIV viremia (29). Therefore, the development of CAR/CXCR5-T cells that are further engineered to be resistant to HIV infection and less susceptible to exhaustion may improve efficacy. Future studies might include strategies to generate HIV-resistant CAR T cells by disruption of the HIV coreceptor CCR5 to prevent HIV infections (60–64) and/or disruption of PD-1 expression or other inhibitory molecules in CAR T cells to prevent exhaustion and improve efficacy (65, 66).

In conclusion, the current study supports the use of the hDRAGA mouse model for HIV immunotherapy studies. Engineered T cells were successfully produced from hDRAGA mouse splenocytes. The CAR/CXCR5-T cells persisted in peripheral blood and secondary lymphoid tissues through the end of the study at 28 DPI. CAR/CXCR5-T cells co-localized with HIV-producing cells in B cell-containing follicle-like structures. However, CAR/CXCR5-T cells did not reduce HIV-1 viral loads in plasma, likely due to the infused cells becoming infected. Additional studies investigating CAR/CXCR5-T cell immunotherapy as a strategy to provide long-term durable virologic control of HIV infection are warranted. Future studies might include generating HIV-resistant and/or exhaustion-resistant CAR/CXCR5-T cells.

## Acknowledgments

S.A.C and M.R. are U.S. Federal employees. The work of these individuals was prepared as part of official government duties. Title 17 U.S. C. §105 provides that “copyright protection under title is not available for any work of the United States Government.” Title 17 U.S.C. §101 defines U.S. Government work as work prepared by a military service member or employee of the U.S. Government as part of that person’s official duties. The views expressed are those of the authors and do not necessarily reflect the official policy or position of the Department of the Navy, Department of the Army, Department of Defense, or the U.S. Government. The following reagent was obtained through the NIH HIV Reagent Program, Division of AIDS, NIAID, NIH: Human Immunodeficiency Virus Type 1 HIV-1_Ba-L_ virus (ARP-510), contributed by Dr. Suzanne Gartner, Dr. Mikulas Popovic, and Dr. Robert Gallo. Anti-CD3 (OKT3) used in the current study was provided by the National Cancer Institute’s Biological Resources Branch Preclinical Repository. We thank James Berg and Ian Gorrell-Brown for sectioned slide preparation and RNAscope imaging. Some figures were created with BioRender.

## Disclaimer

This material has been reviewed by the Walter Reed Army Institute of Research. There is no objection to its presentation and/or publication. The opinions or assertions contained herein are the private views of the author, and are not to be construed as official, or as reflecting true views of the Department of the Army or the Department of Defense.

P.J.S is a cofounder of MarPam Pharma, LLC

## Footnotes

This work was supported in part by grants from the Military Infectious Diseases Research Program to SAC, work unit A1210.

## Abbreviations used in this article

hDRAGA: Humanized DRAGA
CAR: Chimeric Antigen
Receptor DMPA: Medroxyprogesterone
Acetate DPI: Days post-infusion
HSC: Hematopoietic Stem Cells
MBL: mannose-binding lectin
NHP: non-human primate
RMs: Rhesus Macaques
SHIV: Simian/HIV (SHIV) viruses
WRAIR/NMRC: Walter Reed Army Institute of Research/Naval Medical Research Command

