## Supplementary Figure 1 and Supplementary Table I for "Human-immune-system humanized-DRAGA mice are a valuable model to study novel immunotherapies for HIV-1"

### **Supplementary data**

**Supplementary Figure 1. Animal allocation considerations prior to CAR/CXCR5-T cell infusion.** Criteria of control and treated animals included age, the peak of plasma HIV RNA viral loads, levels of plasma HIV RNA viral loads prior to infusion of transduced cell products, weight, human CD4^+^/CD8^+^ T cell ratios, and human CD4^+^ T cell counts. *p* values were determined by the two-tailed Mann–Whitney U test. Each square or circle represents an individual mouse. Red = control (untreated) mice; blue = treated mice. The bar and error represent mean ± SD or geometric mean with 95% CI (for CD4^+^ T cell count).

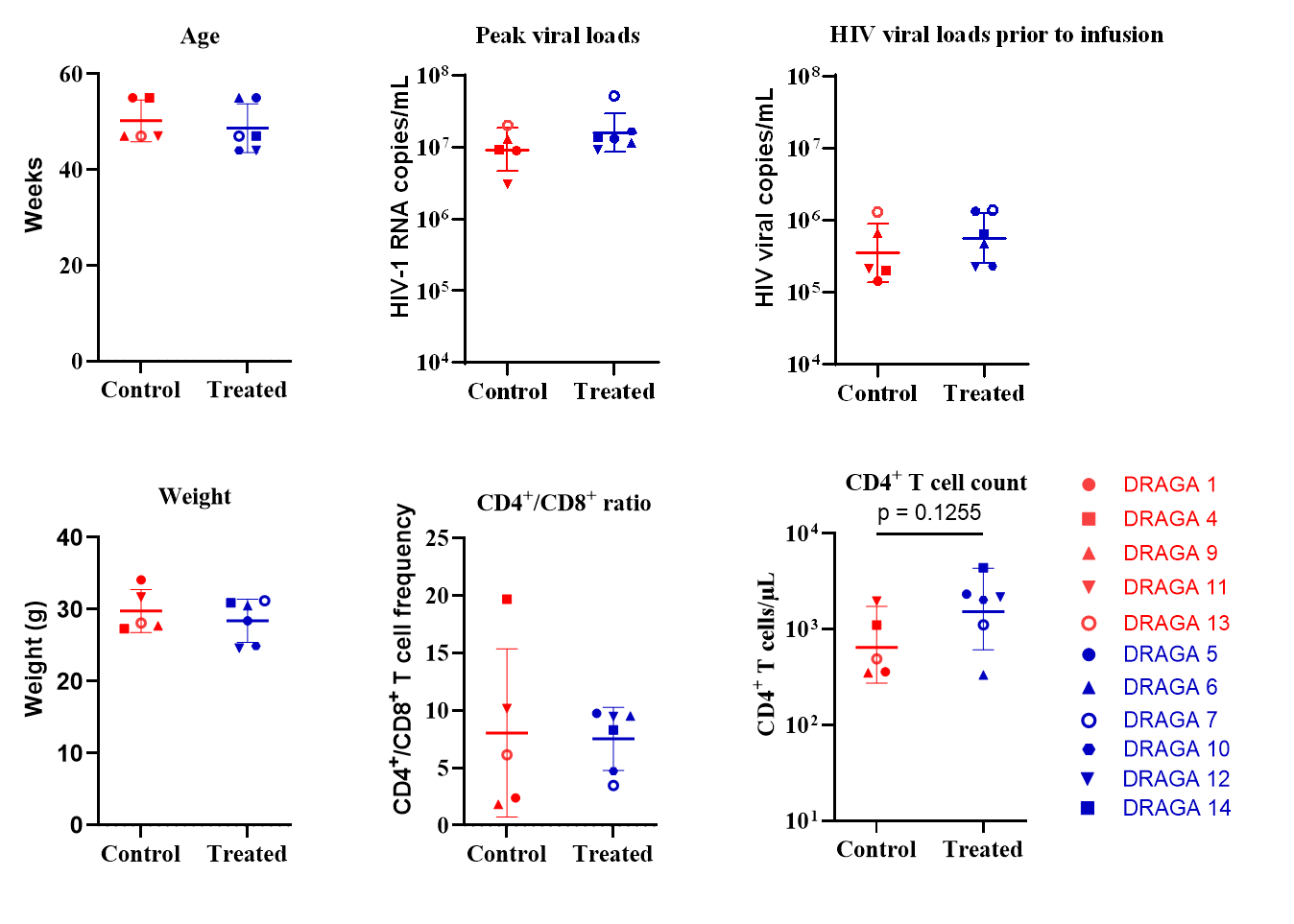

**Supplementary Table I. HIV-1 plasma viral loads in treated and control animals post-cell infusion**

| **Day(s) post-infusion** | **Control animals**  **(x 10^5^ copies/ml)** | | | | | | | **Treated animals**  **(x 10^5^ copies/ml)** | | | | | | | |
| --- | --- | --- | --- | --- | --- | --- | --- | --- | --- | --- | --- | --- | --- | --- | --- |
|  | **1** | **4** | **9** | **11** | **13** | **Mean** | **SD** | **5** | **6** | **7** | **10** | **12** | **14** | **Mean** | **SD** |
| 0 | 1.42 | 2.00 | 6.66 | 2.09 | 13.00 | 5.03 | 4.93 | 13.20 | 4.70 | 13.90 | 2.29 | 2.29 | 6.44 | 7.14 | 5.21 |
| 6 | 11.30 | 11.50 | 21.80 | 13.10 | 43.80 | 20.30 | 13.80 | 30.00 | 17.40 | 20.10 | 19.00 | 12.80 | 11.50 | 18.50 | 6.70 |
| 14 | 13.00 | 5.42 | 6.22 |  |  | 8.21 | 4.16 | 12.80 | 7.96 | 10.40 | 2.19 |  |  | 8.35 | 4.55 |
| 28 | 6.16 |  |  |  |  | 6.16 |  | 33.5 | 14.7 |  |  |  |  | 24.1 | 13.3 |
